# Targeting Specific Kinase Substrates Rescues Increased Colitis Severity Induced by the Crohn’s Disease-Linked LRRK2-N2081D Variant

**DOI:** 10.1101/2024.09.05.611530

**Authors:** George R. Heaton, Xianting Li, Xiaoting Zhou, Yuanxi Zhang, Duc Tung Vu, Marc Oeller, Ozge Karayel, Quyen Q. Hoang, Meltem Ece Kars, Minghui Wang, Leonid Tarassishin, Matthias Mann, Inga Peter, Zhenyu Yue

## Abstract

LRRK2 contains a kinase domain where both the N2081D Crohn’s disease (CD) risk and the G2019S Parkinson’s disease (PD)-pathogenic variants are located. The mechanisms by which the N2081D variant increase CD risk, and how these adjacent mutations result in distinct diseases, remain unclear. To investigate the pathophysiology of the CD-linked LRRK2 N2081D variant, we generated a knock-in (KI) mouse model and compared its effects to those of the LRRK2-G2019S mutation. We find that *Lrrk2^N^*^2081*D*^ KI mice demonstrate heightened sensitivity to induced colitis, resulting in more severe inflammation and intestinal damage than *Lrrk2^G2019S^*KI and wild-type mice. Analysis of Colon tissue revealed distinct mutation-dependent LRRK2 RAB substrate phosphorylation, with significantly elevated phosphorylated RAB10 levels in *Lrrk2^N2081D^* mice. In cells, we demonstrate that the N2081D mutation activates LRRK2 through a mechanism distinct from that of LRRK2-G2019S. We further find that proinflammatory stimulation enhances LRRK2 kinase activity, leading to mutation-dependent differences in RAB phosphorylation and inflammatory responses in dendritic cells. Finally, we show that genetic knockout of *Rab12*, but not pharmacological LRRK2 kinase inhibition, significantly reduced colitis severity in *Lrrk2^N2081D^* mice. Our study characterizes the pathogenic mechanisms of LRRK2-linked CD, highlights important structural and functional differences between disease-associated LRRK2 variants, and suggests RAB proteins as promising therapeutic targets for modulating LRRK2 activity in CD treatment.

## Main

Mutations in *LRRK2* are a major cause of Parkinson’s disease (PD), a common age-related neurodegenerative disorder characterized by the degeneration of dopaminergic neurons in the substantia nigra pars compacta and the presence of aggregated alpha-synuclein in neurons.^1,2^ The *LRRK2* gene encodes Leucine-rich repeat kinase 2 (LRRK2), a large, complex protein with multiple functional regions.^3^ The catalytic core of LRRK2 consists of a GTPase and an adjacent serine/threonine kinase domain. LRRK2 can auto-phosphorylate at serine residue S1292 in its leucine-rich repeat (LRR) region and phosphorylate RAB GTPases at a conserved site within their Switch II domain.^4^ A growing body of evidence suggests that LRRK2 plays a significant role in regulating membrane trafficking at various stages of the endo-lysosomal pathway, including a response to damaged lysosomes through its interaction with and phosphorylation of RAB GTPases.^4–10^

Our previous exome-wide association analysis of Crohn’s disease (CD) in individuals of Ashkenazi Jewish descent identified the N2081D coding variant in LRRK2 as a CD risk allele.^3^ CD is a type of inflammatory bowel disease (IBD), a chronic disorder affecting the gastrointestinal tract.^11^ In our present study, we confirmed the genetic pleiotropy of LRRK2 in both CD and PD through association analysis of three large biobank datasets. To understand the mechanistic basis of these observations—specifically, how the N2081D variant affects LRRK2 function to increase CD risk and its similarities and differences with the PD-associated G2019S variant that might explain their distinct phenotypes—we developed a KI mouse model carrying the LRRK2 N2081D mutation using CRISPR/Cas9 gene editing.

Our findings show that *Lrrk2^N2081D^* KI mice exhibit significantly increased intestinal inflammation and colitis severity, confirming the N2081D variant as a pathogenic risk factor. Comparative tissue analysis from *Lrrk2^G2019S^*and *Lrrk2^N2081D^* mutation carriers further revealed significant differences in LRRK2 RAB substrate phosphorylation, demonstrating distinct functional effects of these variants. Mechanistically, we find that the N2081D variant activates LRRK2 by disrupting an interdomain interaction between the LRR and kinase regions. Furthermore, proinflammatory stimulation enhances LRRK2 kinase activity, leading to mutation-dependent differences in RAB phosphorylation and inflammatory responses.

Finally, we demonstrated that genetic targeting of specific LRRK2 substrates reduces colitis severity in *Lrrk2^N2081D^* mutation carriers, whereas a small molecule LRRK2 inhibitor did not produce a clear therapeutic effect. Our findings highlight distinct pathological features of the LRRK2-N2081D variant and suggest potential therapeutic targets for CD treatment.

### Association Analysis of Crohn’s Disease and Parkinson’s Disease-Linked Variants Confirms Genetic Pleiotropy of LRRK2

We examined the associations of the LRRK2 G2019S and N2081D variants with CD and PD across three biobanks: BioMe BioBank, UK Biobank (Genebass), and the VA Million Veteran Program (MVP).^12,13^ We conducted population-specific association tests and four meta-analyses, including a multi-ancestry meta-analysis that comprised up to 1,072,253 individuals. Our analysis uncovered significant differences in the odds ratios of these LRRK2 variants for CD and PD (Fig. 1a and Supp. Table 1 & 2). Specifically, the N2081D variant increases the risk for both CD and PD, whereas the G2019S variant was significantly associated with an increased risk for PD in all European-specific analyses and meta-analyses. Furthermore, G2019S increased the risk of CD in the European-only analyses of BioMe BioBank, MVP and meta-analysis of MVP. However, its association with CD in UK Biobank was inconsistent with previous reports, as none of the CD cases in UK Biobank carried G2019S variant, likely due to significant differences in G2019S risk allele frequencies in various ancestral groups. Overall, these findings further demonstrate the extent of the genetic pleiotropy between CD and PD within the LRRK2 gene.^3,14^

**Fig. 1.**
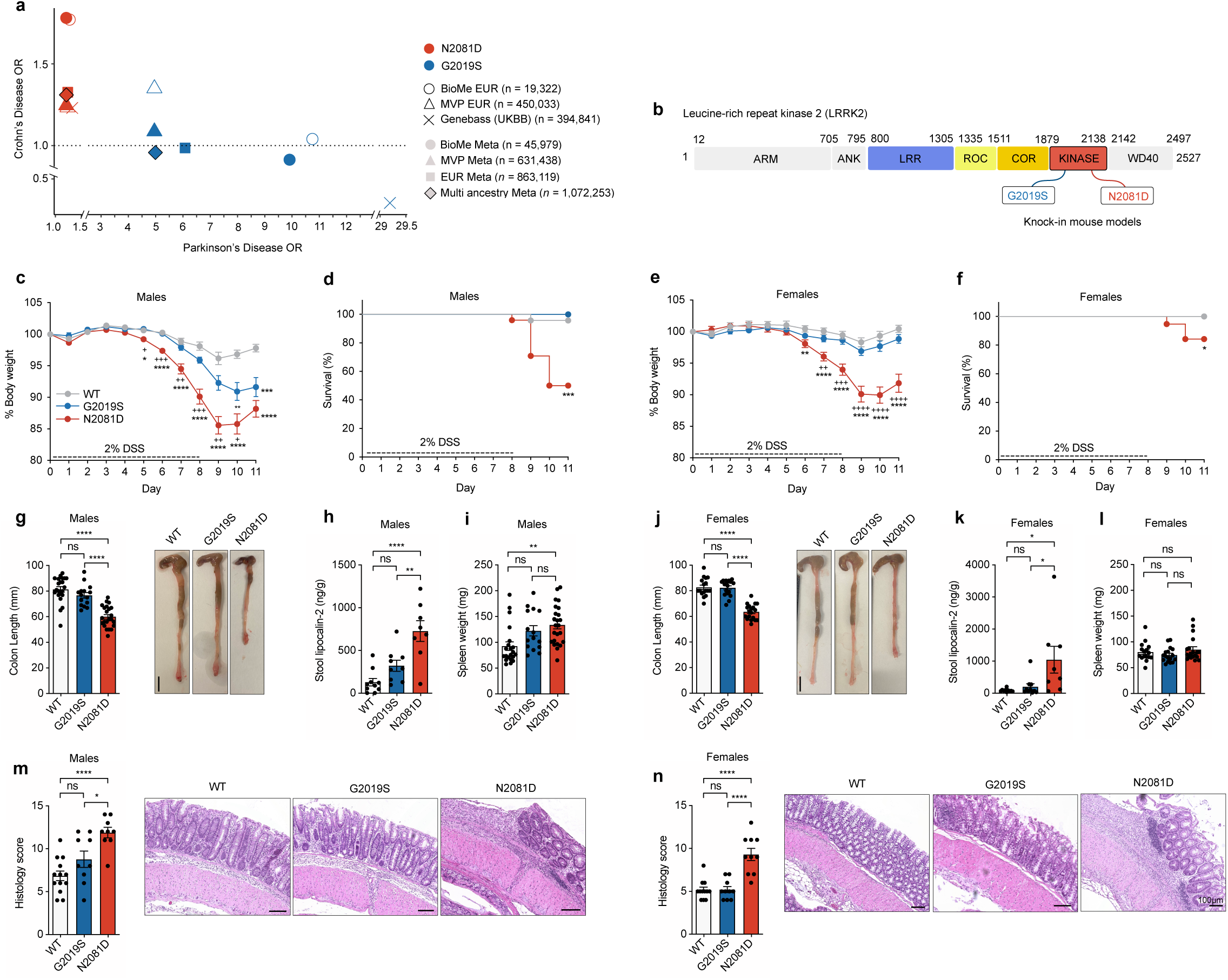
Validation of the LRRK2-N2081D Knock in mouse model of Crohn’s disease. **(a)** Odds ratios (OR) for the associations of the G2019S and N2081D LRRK2 variants with CD and PD. **(b)** Schematic representation of the LRRK2 protein, highlighting the protein domains and the locations of disease-associated variants. **(c, e)** Normalized percentage of body weight through the course of acute DSS treatment for male and female mice, respectively (*n* = 61 male, 53 female animals). * indicates a significant difference between WT and either G2019S or N2081D; + indicates a significant difference between N2081D and G2019S. **(d, f)** Survival curves of male and female mice; animals surpassing humane endpoints were sacrificed (*n* = 61 male, 53 female animals). **(g, j)** Measurements of colon length and representative images for male and female mice, respectively (*n* = 60 male, 53 female animals). **(h, k)** Stool Lipocalin-2 measured by ELISA for males and females (*n* = 27 male, 28 female animals). **(i, l)** Weight of spleen recorded on sacrifice for Males and Females (*n* = 60 male, 52 female animals). **(m, n)** Representative images and histological scoring of intestinal inflammation and damage in male and female mice (*n* = 31 male, 32 female animals).

### *LRRK2^N2081D^* Mice Show Increased Sensitivity to Chemically-Induced Colitis

We developed a KI mouse model of the LRRK2-N2081D variant using CRISPR-Cas9 (**Supp. Fig. 1a**). This model carries the *Lrrk2^N2081D^* murine equivalent of the human N2081D LRRK2 variant, expressed at endogenous levels. Additionally, we used a previously established *Lrrk2* KI model with the G2019S mutation (*Lrrk2^G2019S^*).^15^ Homozygous N2081D KI mice appeared grossly normal compared to Wild-type (WT) and *Lrrk2^G2019S^* mice (**Supp. Fig. 1 b-d**).

To evaluate the impact of LRRK2 mutations on colonic inflammation and colitis symptoms, we employed the widely used DSS model. Prolonged DSS exposure leads to epithelial cell death and compromised intestinal barrier integrity, resulting in leukocyte infiltration and colonic tissue damage.^16^ We induced colitis by administering 2% DSS in the drinking water to 16-week-old male and female WT mice and homozygous carriers of the *Lrrk2^G2019S^* or *Lrrk2^N2081D^* variant for 8 days, followed by 3 days of regular water for partial recovery.

In both sexes, *Lrrk2^N2081D^* mice exhibited more severe weight loss compared to WT and *Lrrk2^G2019S^* mice (**Fig. 1c & e**). Male mice experienced the most severe weight loss, with a significant number of N2081D males requiring early sacrifice due to the severity of colitis symptoms (**Fig. 1d**). At the experimental end point, both male and female *Lrrk2^N2081D^* mice had shorter colons than their sex-matched WT and *Lrrk2^G2019S^* counterparts, indicating a greater degree of colonic inflammation (**Fig. 1g & j**). Consistent with these results, stool samples extracted from colons of N2081D mutation carriers had significantly higher levels of the proinflammatory marker lipocalin-2 (**Fig. 1h & k**). Male N2081D carriers also had increased spleen weight compared to WT mice, suggesting a heighted immune response (**Fig. 1l**).

Histological examination of colon samples revealed a greater degree of inflammation and tissue damage in both male and female *Lrrk2^N2081D^* carriers compared to sex-matched *Lrrk2^G2019S^* and WT mice (**Fig. 1m & n)**. Collectively, these results show that the *Lrrk2^N2081D^* mutation significantly exacerbates colonic inflammation and the severity of induced colitis, validating the *Lrrk2^N2081D^* model as a tool to study LRRK2-linked CD pathogenic mechanisms.

### Distinct Functional Effects of Crohn’s Disease and Parkinson’s Disease-Linked LRRK2 Variants

To assess the effects of the G2019S and N2081D mutations on LRRK2 kinase activity, we first measured levels of established LRRK2-kinase targets in brain and colon tissues of KI mouse models. Analysis of ∼3 month old male mice revealed elevated levels of the phosphorylated LRRK2-S1292 in *Lrrk2^G2019S^* mutation carriers with significant differences compared to WT and *Lrrk2^N2081D^* animals observed in the brain. Notably, increased phosphorylation of RAB12 was exclusively observed in *Lrrk2^G2019S^*mice in both brain and colon tissue. However, LRRK2-G2019S only modestly increased RAB10 phosphorylation levels compared to WT (**Fig 2a & b**).

**Fig. 2.**
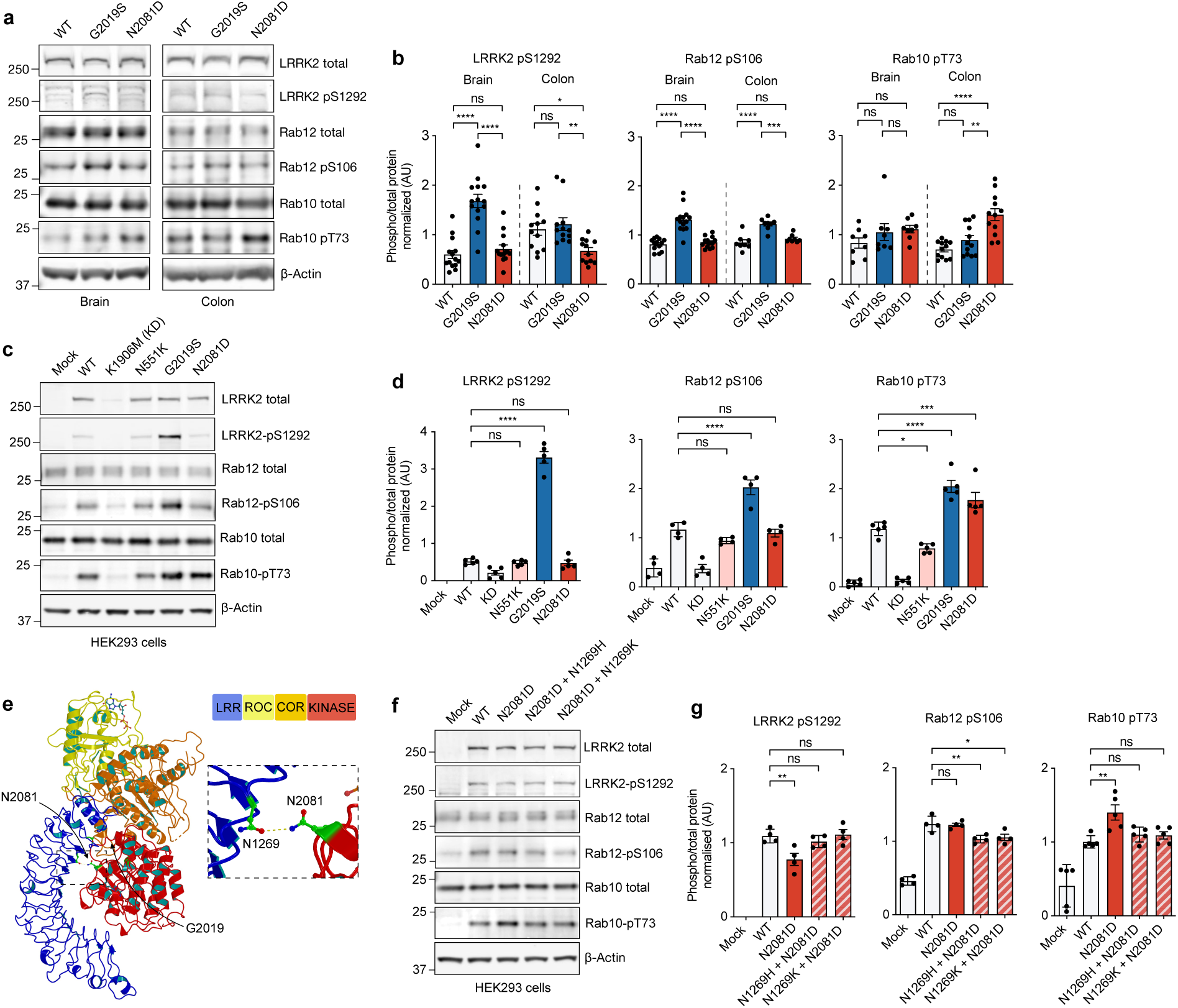
Kinase activity characterization of Parkinson’s disease and Crohn’s disease linked LRRK2 variants. **(a)** Representative immunoblots of Colon and Brain tissue lysates of WT, G2019S and N2081D knock-in animals. **(b)** Quantification of endogenous LRRK2-S1292 autophosphorylation and Rab substrate phosphorylation (*n* = 42 animals). **(c)** Representative immunoblots of HEK293 cells following expression of LRRK2 variants. **(d)** Quantification of LRRK2 S1292 autophosphorylation and phosphorylation of endogenous RAB substrates (*n* = 5 independent experiments). **(e)** Structural modeling of LRRK2 domains (LRR, ROC, COR, and Kinase) based on the full-length structure, with the positions of N2081 and G2019 residues highlighted. **(f)** Representative immunoblots of HEK293 cells expressing LRRK2 mutants at N2081 and N1269 residues. **(g)** Quantification of LRRK2 autophosphorylation and endogenous RAB phosphorylation (*n* = 5 independent experiments).

In contrast, N2081D mutation carriers demonstrate significantly higher levels of phosphorylated RAB10 in colon tissues, with a similar trend in the brain, compared to WT and G2019S mice (**Fig 2a & b**). A comparison of brain and colon samples from N2081D mutation carriers showed significantly higher levels of phosphorylated RAB10 in the colon than in the brain, which may suggest that this variant has a stronger effect in this tissue (**Supp. Fig 2a & b**). Analysis of a subset of N2081D male mice with severe colitis symptoms showed a striking increase in phosphorylated RAB10 in DSS-treated colon tissue compared to untreated animals, indicating that the N2081D mutation’s effects may be further amplified by severe inflammation **(Supp. Fig. 2c & d)**. Overall, our findings suggest that while both the N2081D and G2019S mutations enhance LRRK2 kinase activity, their distinct patterns of substrate phosphorylation indicate different mechanisms of action.

To further investigate the functional effects of CD- and PD-linked LRRK2 variants, we assessed their impact on LRRK2 kinase activity in HEK293 cells (**Fig 2c & d)**. Mock transfection and the kinase-dead (KD) K1906M variant served as negative controls. Consistent with our observations in mouse tissue, only LRRK2-G2019S significantly enhanced LRRK2-S1292 autophosphorylation. Increased RAB12-S106 phosphorylation was also observed only following LRRK2-G2019S expression. Both the N2081D and G2019S variants induced significant increases in RAB10 phosphorylation, while the CD-protective N551K variant decreased RAB10 phosphorylation levels (**Fig. 2c & d).**

To investigate the mechanism of the N2081D mutation, we examined previously reported high-resolution structural models of inactive full-length LRRK2 (**Fig. 2e**). These models predict a hydrophilic interaction between the kinase domain’s N2081 residue and the N1269 residue in the LRR domain. In its inactive state, the LRR domain tightly interacts with the kinase domain, blocking the active site. The N2081D mutation is expected to disrupt this interaction, exposing the kinase domain and allowing greater access to RAB substrates. To validate this proposed mechanism, we introduced mutations at the N1269 residue to re-establish its interaction with the N2081D mutant and evaluated the impact on LRRK2’s kinase activity. Our data show that the N1269H and N1269K mutations effectively counteract N2081D-mediated activation of LRRK2, reducing RAB10 phosphorylation to near WT levels (**Fig. 2f & g**). Although LRRK2-N2081D alone did not increase RAB12 phosphorylation, the compensatory N1269 mutations also decreased RAB12 phosphorylation relative to WT. Additionally, the N2081D variant caused a modest decrease in S1292 phosphorylation compared to WT. This reduction likely results from the disruption of the N1269-N2081 interaction, causing separation between the S1292 residue in the LRR domain and the kinase domain’s active site. Supporting this, the N1269 mutations restored S1292 phosphorylation to WT levels in the presence of the N2081D variant (**Fig. 2f & g)**.

In summary, our findings suggest that the N2081D mutation activates LRRK2 by inducing a conformational shift, resulting in functional effects distinct from the PD-associated LRRK2 G2019S variant.

### Dendritic Cell Stimulation Induces LRRK2 Mutation-Dependent Differences in RAB Phosphorylation and Inflammatory Responses

Dendritic cells are key mediators of DSS-induced colitis by initiating and amplifying the inflammatory response.^17–19^ Several studies have also demonstrated a critical role for LRRK2 in dendritic cell biology.^7,18,20,21^ To investigate the impact of each LRRK2 mutation in primary cells, we isolated bone marrow-derived dendritic cells (BMDCs) from WT, *Lrrk2^G2019S^* and *Lrrk2^N2081D^* mice. In untreated BMDCs, functional effects of *Lrrk2^G2019S^* and *Lrrk2^N2081D^* mirrored those seen in HEK293 cells, with increased phosphorylation of S1292, RAB12, and RAB10 in *Lrrk2^G2019S^* compared to WT, while the *Lrrk2^N2081D^* variant specifically increased RAB10 phosphorylation (**Fig. 3a**). These results indicate that the activation mechanisms identified in HEK293 cells are conserved in primary dendritic cells from LRRK2 KI mouse models.

**Fig. 3.**
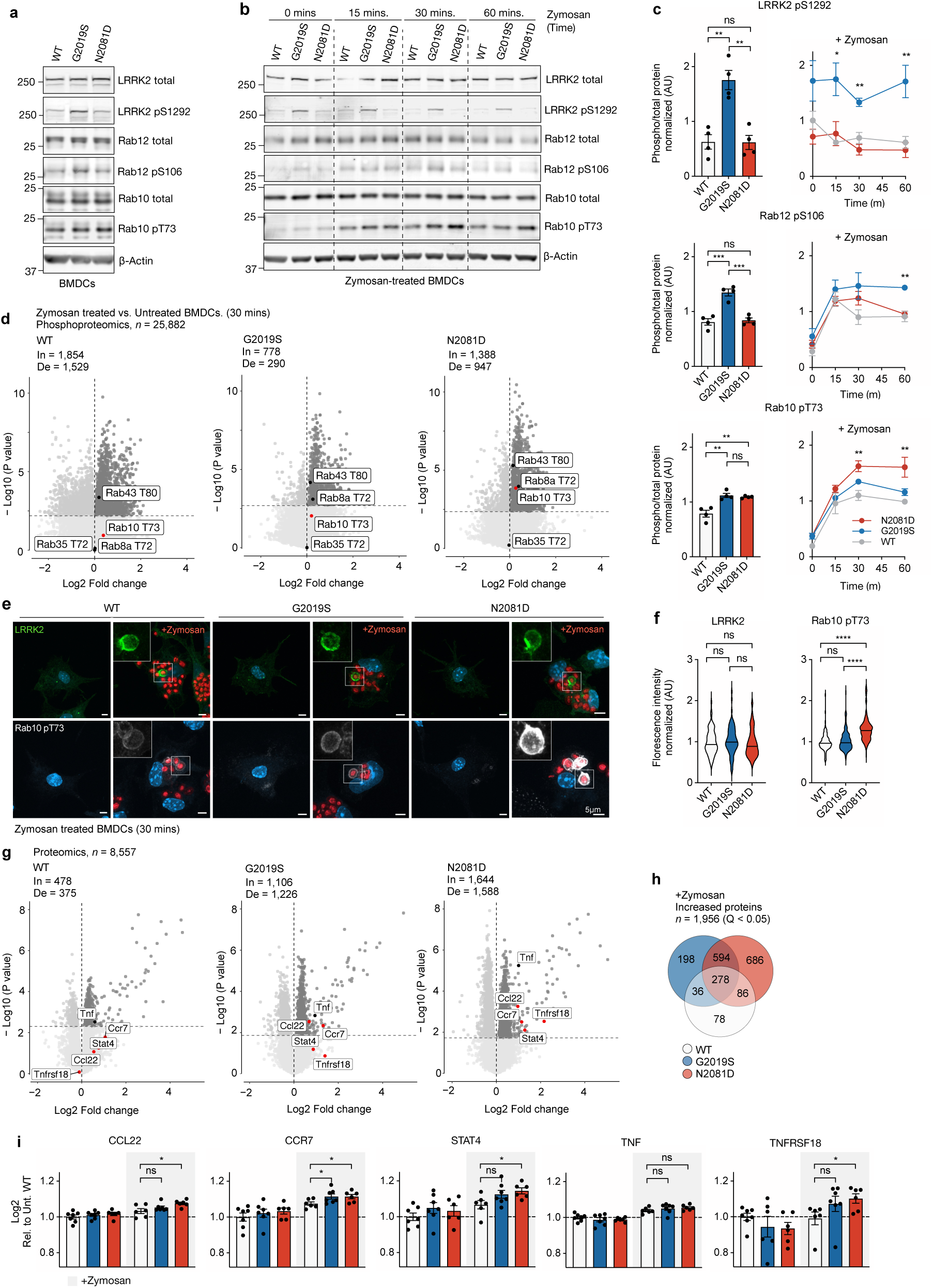
Activation of Dendritic Cells Induces LRRK2 Mutation-Dependent Differences in RAB Phosphorylation and Inflammatory Signaling. **(a)** Representative immunoblot of untreated WT, G2019S, and N2081D BMDCs. (*n* = 4 individual experiments) **(b)** Representative immunoblot of WT, G2019S, and N2081D BMDCs treated with zymosan for 60 minutes (*n* = 4 individual experiments). **(c)** Immunoblot quantification of untreated and zymosan-treated cells. **(d)** Phosphoproteomics volcano plot of WT, G2019S, and N2081D BMDCs treated with Zymosan for 30 minutes. Established LRRK2- phosphorylated RABs are annotated. The horizontal dashed line represents a q-value threshold of <0.05. "In" and "De" indicate the number of proteins with increased or decreased phosphorylation surpassing this threshold, respectively. (*n* = 24 animals used for BMDC cultures, 8 individual experiments). **(e)** Representative images of endogenous LRRK2 and phosphorylated RAB10 following 30 minutes of zymosan treatment. **(f)** Quantification of LRRK2 and RAB10 pT73 fluorescence intensity on the surface of ingested zymosan (LRRK2, *n* = 216 images, 4 independent experiments; pRAB10, *n* = 281 images, 5 independent experiments). **(g)** Proteomics volcano plot of WT, G2019S, and N2081D BMDCs treated with zymosan for 30 minutes. **(h)** Venn diagram showing significantly increased proteins following Zymosan treatment. **(i)** Log2-transformed proteomics data normalized to untreated WT cells (*P < 0.05 by two-tailed t-test).

Given the increased RAB10 phosphorylation observed in *Lrrk2^N2081D^*mutation carriers after severe colonic inflammation, we hypothesized that inflammatory stimuli might activate LRRK2 kinase activity. To test this, BMDCs were stimulated with zymosan, a proinflammatory agent that activates dendritic cells through pattern recognition receptors. Immunoblot analysis confirmed activation of the NF-κB pathway across all genotypes. Interestingly, no significant differences were observed in IκBα and IKKβ degradation or in the phosphorylation of NF-κB p65 between genotypes **(Supp. Fig. 3)**. Following zymosan treatment, we observed a rapid and striking increase in RAB phosphorylation across all genotypes, but with clear LRRK2 mutation-dependent differences in effect size. *Lrrk2^G2019S^* cells exhibited a progressive rise in RAB12 phosphorylation, peaking higher than in WT and N2081D cells. Conversely, N2081D cells showed a significant increase in RAB10 phosphorylation relative to G2019S and WT. This finding indicates that dendritic cell stimulation activates LRRK2 kinase activity, resulting in mutation-dependent differences in substrate phosphorylation (**Fig. 3b & c**).

To further characterize the activation response, we performed mass spectrometry-based phosphoproteomics and proteomics on dendritic cells. Phosphoproteome analysis showed increased phosphorylation of known LRRK2 RAB substrates, including RAB43, RAB8A, and RAB10, following zymosan treatment, while RAB35 phosphorylation remained unchanged. The total levels of these RAB substrate proteins were consistent across all genotypes. In line with our immunoblot data, RAB10 phosphorylation was significantly higher in N2081D cells after zymosan treatment compared to WT cells **(Fig. 3d).** To confirm these findings and explore their mechanistic basis, BMDCs were treated with fluorescently labeled zymosan bioparticles and imaged using confocal microscopy. Zymosan phagocytosis by dendritic cells resulted in the recruitment of LRRK2 to the phagosome surface. Quantification of phosphorylated RAB10 intensity revealed significantly higher accumulation on phagosome surfaces in *Lrrk2^N2081D^*cells compared to WT and *Lrrk2^G2019S^* cells **(Fig. 3e-f)**.

Further analysis of the total proteomics data revealed only minor changes in untreated G2019S and N2081D BMDC cells compared to WT, with very few proteins showing statistically significant differences in abundance (Q < 0.05). Similar results were observed in brain and colon tissue **(Supp. Fig. 4)**. However, Zymosan stimulation in BMDCs resulted in a significant increase in the levels of numerous proteins **(Fig. 3h)**. Among the 1,644 proteins significantly upregulated following Zymosan treatment in N2081D cells (Q < 0.05), several proteins associated with inflammatory signaling were also significantly elevated compared to WT-Zymosan-treated cells. **(Fig. 3i)**. Specifically, we observed increased levels of CCL22, a chemokine involved in recruiting regulatory T cells, and CCR7, a receptor crucial for dendritic cell and T cell migration. Additionally, STAT4, a transcription factor essential for Th1 cell differentiation, and TNFSF18, a member of the TNF superfamily involved in T cell co-stimulation, were elevated. A similar trend was observed in *Lrrk2^G2019S^* cells, though only CCR7 showed a statistically significant increase compared to WT.

### LRRK2 Kinase Inhibitor MLi-2 Does Not Improve Colitis in *Lrrk2^N2081D^* Mice

We next evaluated potential therapeutic strategies to reduce colitis severity in N2081D mutation carriers, focusing on *Lrrk2^N2081D^*male mice, which exhibited the most severe colitis symptoms among our experimental groups. We hypothesized that kinase inhibitor treatment might mitigate the effects of the N2081D mutation and improve symptoms of DSS-induced colitis. To test this, we provided male *Lrrk2^N2081D^* mutation carriers with chow supplemented with MLi-2, a potent and selective LRRK2 inhibitor, targeting a 60 mg/kg/day dosage, as previously described by Kluss et al.^22,23^

*Lrrk2^N2081D^* carriers were provided with facility chow or MLi-2-supplemented chow for 3 days prior to DSS treatment. On day 4, we began administering 2% DSS for 8 days, followed by a 3-day recovery period, with mice continuing their respective diets throughout (**Supp. Fig. 5a & b**). Colon tissue samples analyzed at the experimental endpoint showed significant reductions in RAB10 and RAB12 phosphorylation following MLi-2 treatment, confirming effective target engagement (**Supp. Fig. 5c & d)**. However, MLi-2 had minimal impact on body weight loss, spleen weight, colon length, damage, and inflammation, suggesting that in-diet dosing of the LRRK2 kinase inhibitor MLi-2 does not provide significant protection from DSS-induced colitis in N2081D mutation carriers (**Supp. Fig. 5e-i**).

### Rab12 Knockout Reduces RAB10 Phosphorylation and Alleviates Colitis Severity in *Lrrk2^N2081D^* Mice

Research from our lab and others has identified RAB12 as a key regulator of LRRK2-dependent RAB10 phosphorylation.^24–26^ Notably, we recently demonstrated that Rab12 knockout (KO) prevents LRRK2-G2019S-mediated cilia defects in the mouse brain.^26^ While *Rab12* KO mice exhibit normal growth and reach adulthood, Rab10 KO mice do not survive past embryonic development (data not shown). We hypothesized that targeting *Rab12* through genetic deletion might mitigate the effects of the N2081D mutation on RAB10 phosphorylation and limit its pathogenic impact. To test this, we crossed *Lrrk2^N2081D^*mice with *Rab12* KO mice. *Lrrk2^N2081D^;Rab12^-/-^* mice appeared grossly normal, with no significant differences in body weight, colon length, or spleen size compared to WT, *Lrrk2^N2081D^*, or *Rab12^-/-^*, suggesting that loss of *Rab12* is well tolerated (**Supp. Fig. 6**).

*Lrrk2^N2081D^;Rab12^-/-^* mice showed a significant reduction in phosphorylated RAB10 in colon tissue compared to *Lrrk2^N2081D^;Rab12^+/+^* mice, with only a minor observed effect in the brain (**Fig. 4a & b**). Reduced RAB10 phosphorylation were also detected in BMDCs of *Lrrk2^N2081D^;Rab12^-/-^*animals compared to *Lrrk2^N2081D^* carriers expressing RAB12 (**Fig. 4c**). Furthermore, *Rab12* KO significantly decreased zymosan-induced N2081D-mediated RAB10 phosphorylation, resulting in a reduced peak signal over 60 minutes (**Fig. 4d & e**). These findings demonstrate that Rab12 knockout effectively reduces LRRK2-N2081D-mediated RAB10 phosphorylation.

**Fig. 4.**
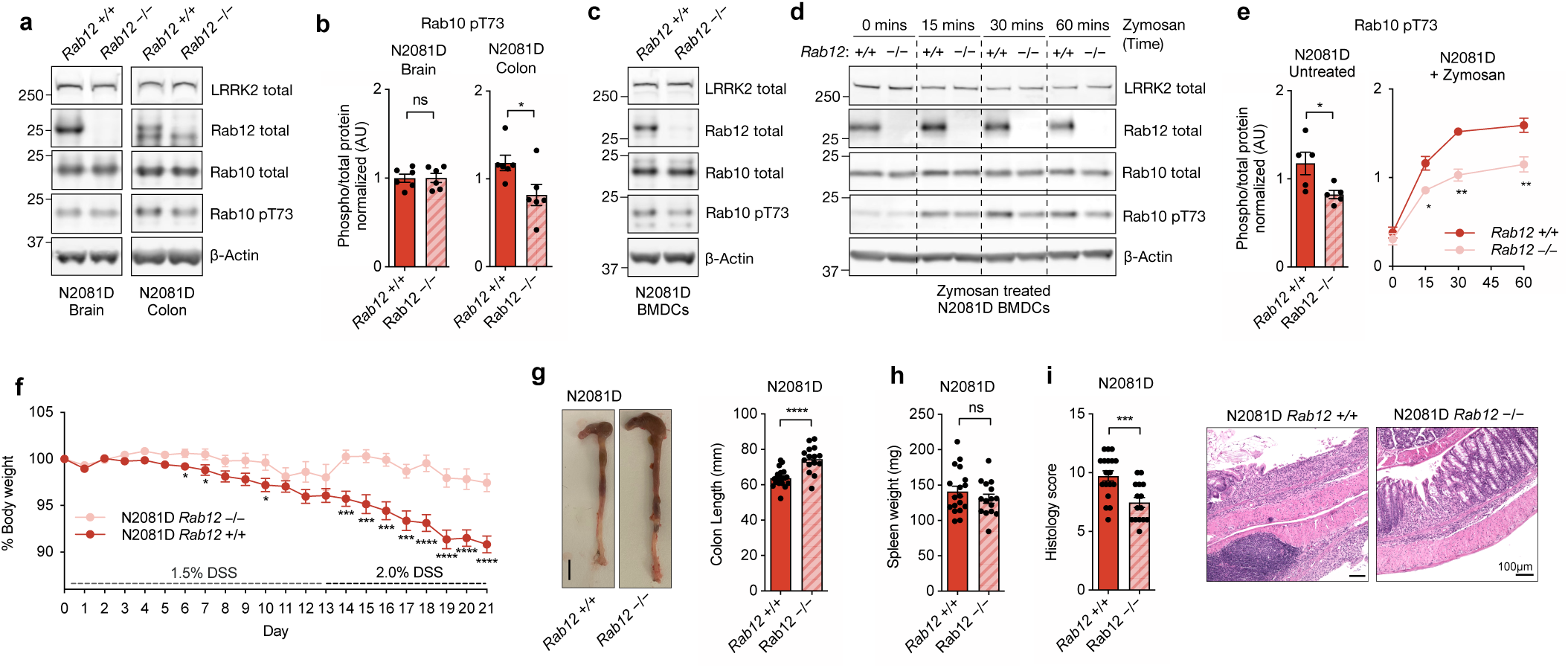
RAB12 Knockout Rescues LRRK2-N2081D Phenotypes. **(a, b)** Representative immunoblot and quantification of phosphorylated RAB10 in the brain and colon from N2081D and N2081D/RAB12 KO mice (*n* = 10 animals). **(c, d)** Representative immunoblot of N2081D and N2081D/RAB12 KO untreated BMDCs (*n* = 10 animals) and following zymosan treatment over the course of 60 minutes (*n* = 8 animals, *n* = 4 independent experiments). **(e)** Immunoblot quantification of untreated and zymosan-treated N2081D and N2081D/RAB12 KO BMDCs. **(f)** Normalized body weight of N2081D and N2081D/RAB12 KO animals during progressively increasing DSS treatment over 21 days. **(g)** Representative colon photos and colon length measurements at sacrifice. **(h)** Spleen weight of N2081D and N2081D/RAB12 KO animals at sacrifice. **(i)** Representative images and histological scoring for colon inflammation and damage (*n* = 33 animals).

We next examined the impact of RAB12 loss on symptoms of induced colitis in *Lrrk2^N2081D^* mice. *Lrrk2^N2081D^;Rab12^-/-^* animals showed significantly less severe DSS-induced body weight loss over the course of DSS-administration (**Fig. 4f**). At the experimental end point, *Lrrk2^N2081D^;Rab12^-/-^*animals had significantly longer colons, indicating reduced colonic inflammation relative to *Lrrk2^N2081D^;Rab12^+/+^* **(Fig. 4g)**. Histological analysis confirmed a lesser degree of colonic damage and immune cell infiltration in RAB12 KO mice. A modest reduction in spleen weight was also observed in *Lrrk2^N2081D^;Rab12^-/-^*mice, although this was not statistically significant **(Fig. 4h & i)**.

In summary, we find that deletion of *Rab12* significantly reduces N2081D-associated colitis severity, highlighting its potential as a therapeutic target. Notably, while WT mice experienced less severe DSS symptoms under the same DSS-dosing regimen, they did not show a measurable benefit from *Rab12* KO. This may suggest that *Rab12* deletion may have a greater therapeutic benefit specifically in LRRK2-mutation carriers (**Supp. Fig. 7**).

## Discussion

Our association analysis of CD- and PD-linked LRRK2 variants across multiple biobank datasets underscores the genetic pleiotropy of the LRRK2 gene. Characterizing mice carrying the LRRK2 N2081D risk variant, we found that it exacerbates induced colitis, confirming its pathogenic role in CD. We further demonstrate that *Rab12* deletion mitigates the pathogenic effects of LRRK2-N2081D, leading to a significant reduction in intestinal inflammation. These findings establish a LRRK2-linked CD mouse model and identify the RAB12-LRRK2 pathway as a promising therapeutic target for modulating LRRK2 activity in CD treatment.

Our study also reveals that, unlike LRRK2-G2019S, the N2081D variant specifically enhances the phosphorylation of RAB10 but not RAB12, whereas the G2019S mutation increases phosphorylation of RAB12 and LRRK2-S1292, with a more modest effect on RAB10 in mouse models. These functional differences likely contribute to the diverse phenotypes observed in LRRK2 mutation carriers, leading to varying degrees of IBD symptoms and PD-associated neurodegeneration in patients.

Structurally, the G2019S substitution replaces glycine with serine within the DYG motif of the activation loop in the active site of the kinase domain, increasing its kinetics without significantly altering the overall conformation.^17,18^ In contrast, our results suggest that the N2081D mutation activates LRRK2 by disrupting an interdomain interaction between the LRR and kinase domains. This conformational shift is crucial for increasing RAB10 phosphorylation, likely by increasing exposure of the kinase active site, but appears to have little effect on the phosphorylation of RAB12. The reasons for this selective effect are not fully understood; however, recent structural analysis suggest that RAB12 and RAB10 bind to distinct sites within the armadillo (ARM) domain of LRRK2.^24,26^ It is possible that the N2081D mutation induces a conformation that preferentially enhances phosphorylation of RABs bound at specific sites on LRRK2 (i.e. RAB10), although this hypothesis requires experimental validation.

In LRRK2-overexpressing HEK293 cells and BMDCs, both N2081D and G2019S variants similarly increased RAB10 phosphorylation. However, zymosan stimulation led to a greater increase in phosphorylated RAB10 in LRRK2-N2081D dendritic cells compared to wild-type and LRRK2-G2019S cells. Elevated RAB10 phosphorylation was also observed in colon tissues of N2081D mutation carriers versus wild-type and G2019S KI animals, with further increases seen in carriers experiencing severe DSS symptoms. Proteomic analysis suggests that elevated RAB10 phosphorylation in LRRK2-N2081D cells coincides with increased inflammatory signaling. This is evidenced by higher levels of proteins associated with immune cell recruitment and activation compared to WT cells following zymosan treatment. Although further confirmation is needed, we propose that these cellular phenotypes likely contribute to the increased severity of colitis observed in LRRK2-N2081D mutation carriers by enhancing immune cell infiltration and sustaining inflammatory responses in intestinal tissue. The precise role of increased RAB10 phosphorylation in this process remains to be determined.

A surprising finding in our study is that dietary administration of the LRRK2 kinase inhibitor MLi-2 did not significantly alleviate DSS-induced colitis in N2081D mice, despite effective target engagement, as evidenced by reduced RAB phosphorylation in colon tissue. Notably, a partial rescue of DSS-colitis severity has been previously observed in *Lrrk2^G2019S^* mice using a similar strategy of dietary MLi-2 dosing.^27^ Additionally, LRRK2 inhibitors have been shown to reduce zymosan-evoked increases in TNF-α and other proinflammatory cytokines in the supernatant of mouse and human cell cultures.^18^ These findings may suggest that an alternative experimental setup or kinase inhibitor dosing strategy could potentially yield a stronger therapeutic effect. However, our current data indicates that kinase inhibition alone may not be sufficient to counteract the heightened sensitivity to induced colitis caused by the N2081D mutation.

In contrast, our study showed that *Rab12* deletion significantly reduced phosphorylated RAB10 levels in colon tissue and led to a significant improvement in colitis severity across multiple readouts. We also found that *Rab12* knockout lowered levels of phosphorylated RAB10 in colon tissue and BMDCs, both under basal conditions and following Zymosan-induced activation. Although the N2081D variant does not increase RAB12 phosphorylation beyond WT levels, our data indicate that N2081D-induced RAB10 hyperphosphorylation depends on the presence of RAB12 and likely its interaction with LRRK2.^26^ While the mechanisms by which RAB12 KO regulates LRRK2 function are still being resolved, recent studies in cellular models have shown that RAB12 depletion disrupts LRRK2-induced perinuclear clustering of lysosomes and impairs LRRK2’s recruitment to damaged lysosomal membranes. Our recent work also demonstrated that the deletion of RAB12 can reverse the reduced cilia formation and increased incidence of split centrosomes caused by LRRK2-G2019S in astrocytes.^26^

Based on our findings, we propose that RAB12 depletion could significantly mitigate the pathogenic effects of the LRRK2 N2081D mutation *in vivo*. However, our understanding of how RAB12 loss and the resulting reduction in RAB10 phosphorylation affect LRRK2-mediated inflammatory pathways remains incomplete. Our results highlight this as a critical area for further investigation.

**Supp. Fig. 1.**
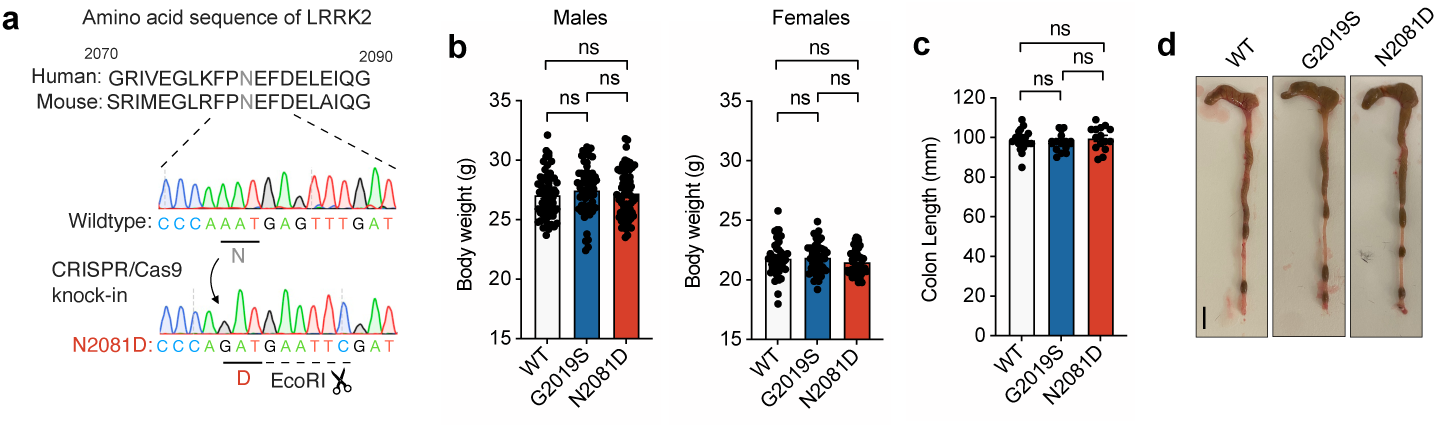
Generation and phenotypic evaluation of LRRK2-N2081D mice. **(a)** Human and mouse amino acid sequence of the LRRK2 kinase domain, with nucleotide sequencing of wildtype and N2081D knockin animals, a silent EcoRI diagnostic site was introduced to identify the N2081D allele. **(b)** Body weight recordings of 16 week old male and female mice (*n* = 180 males, 130 females). **(c)** Measurement of colons from WT, G2019S KI, and N2081D KI 16-week old male mice. **(d)** Representative photograph of colons from the same groups (*n* = 50 males).

**Supp. Fig. 2.**
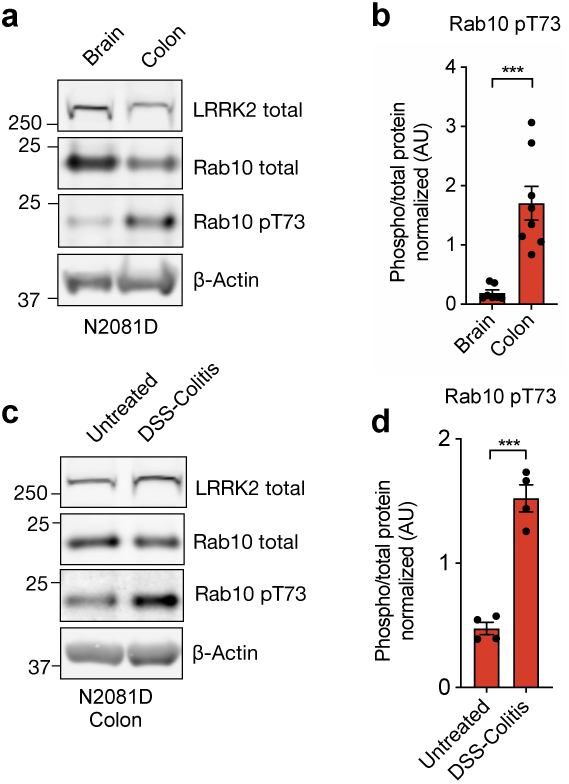
Tissue analysis of RAB10 Phosphorylation in LRRK2-N2081D Mutation Carriers. **(a, b)** Representative immunoblot and quantification of phosphorylated RAB10 in brain and colon tissue of LRRK2-N2081D animals (*n* = 16 animals). **(c, d)** Representative immunoblot and quantification of phosphorylated RAB10 in colon samples from untreated LRRK2-N2081D mice and LRRK2-N2081D mice with severe colitis symptoms (*n* = 8 animals).

**Supp. Fig. 3.**
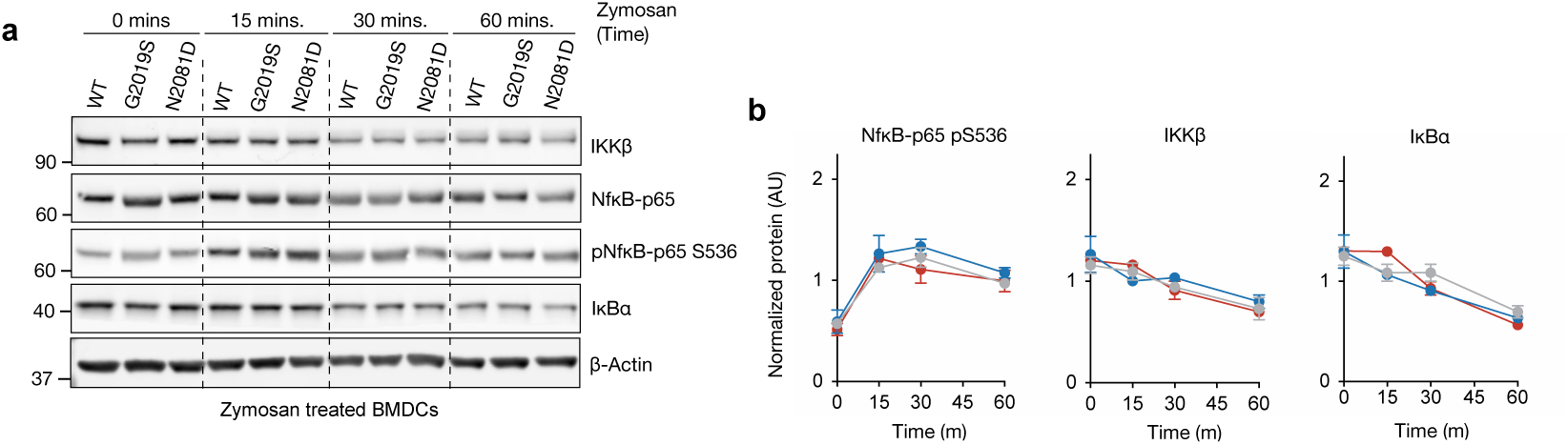
Zymosan Treatment Activates the NF-κB Pathway. **(a)** Representative immunoblot of WT, G2019S, and N2081D BMDCs treated with zymosan over 60 minutes (*n* = 12 animals, 4 individual experiments). **(b)** Immunoblot quantification of zymosan-treated BMDCs.

**Supp. Fig. 4.**
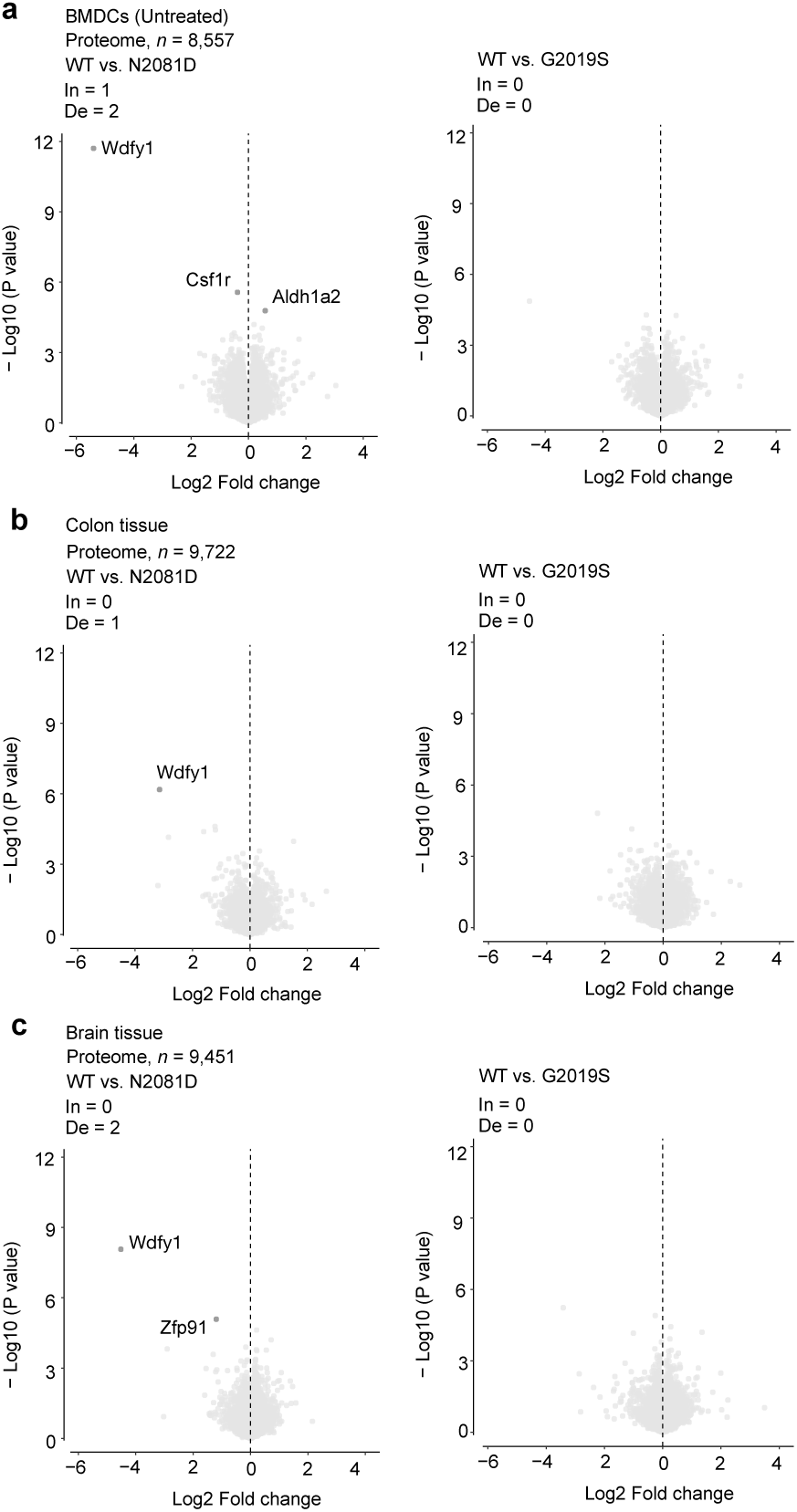
Proteomics Volcano Plots of WT, G2019S, and N2081D Samples. (a) Volcano plot comparing untreated BMDCs from G2019S and N2081D mice to WT. (b) Volcano plot comparing colon tissue from G2019S and N2081D mice to WT. (c) Volcano plot comparing brain tissue from G2019S and N2081D mice to WT (*n* = 24 animals, with BMDCs, colon, and brain tissues collected from the same animals).

**Supp. Fig. 5.**
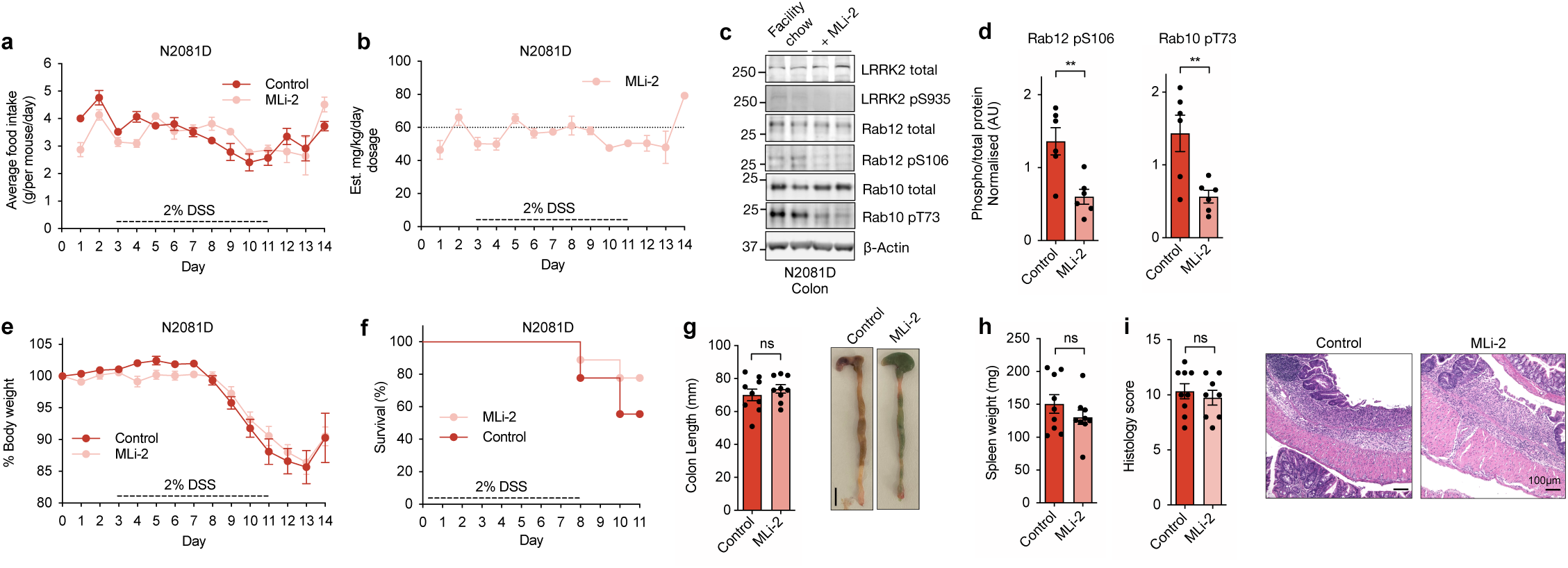
Treatment of LRRK2-N2081D Animals with LRRK2 Kinase Inhibitor MLi-2 Supplemented Chow. **(a)** Average food intake per cage of N2081D mice on facility chow or MLi-2 supplemented chow. **(b)** Estimated mg/kg/day dosage of MLi-2. **(c, d)** Representative immunoblot of colon tissue of N2081D mice and quantification of LRRK2 RAB substrates. **(e)** Normalized percentage of body weight during DSS treatment. **(f)** Survival curves for LRRK2-N2081D males; animals surpassing humane endpoints were sacrificed. **(g)** Representative images and recorded length of colon at sacrifice. **(h)** Spleen weight recorded at sacrifice. **(i)** Representative histology images and scoring (*n* = 17 animals).

**Supp. Fig. 6.**
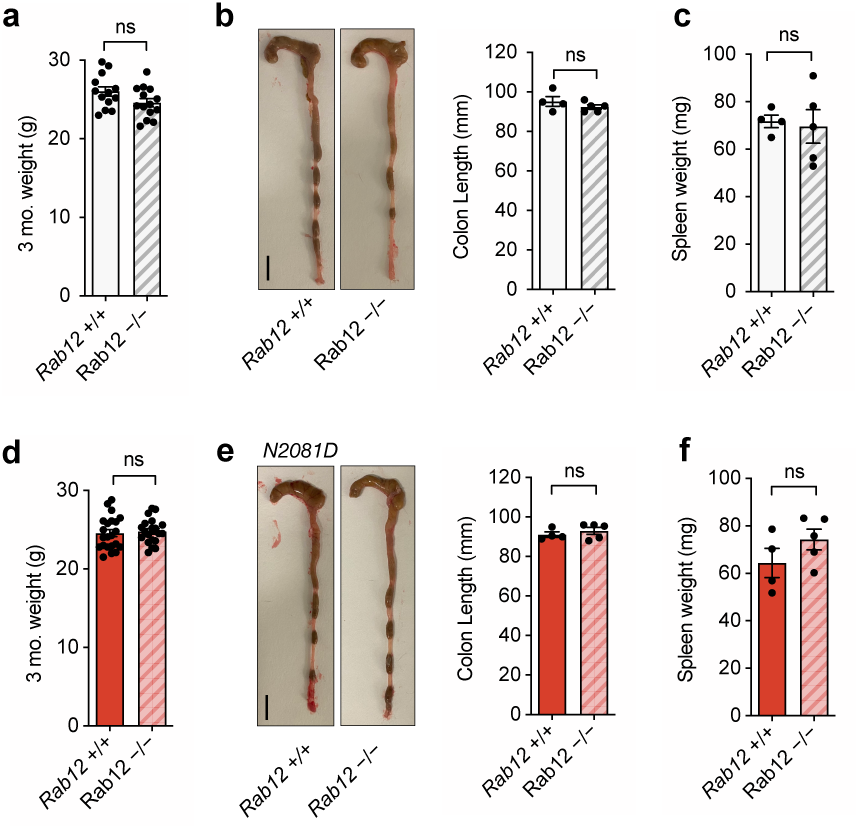
Assessment of RAB12 Knockout on body weight, colon Length and spleen weight. **(a)** Body weight recordings at 3 months of age in WT and RAB12 KO male animals (n = 27). **(b)** Photos of colons and measurements of colon length in WT and RAB12 KO male animals (n = 9). (c) Spleen weight recordings of WT and RAB12 KO male animals (n = 9). **(d)** Body weight recordings at 3 months of age in N2081D and N2081D/RAB12 KO male mice (*n* = 42). **(e)** Photos of colons and measurements of colon length in N2081D and N2081D-RAB12 KO male animals (n = 9). **(f)** Spleen weight recordings of N2081D and N2081D-RAB12 KO male animals (n = 9). All samples were taken from mice that were 3 months old.

**Supp. Fig. 7.**
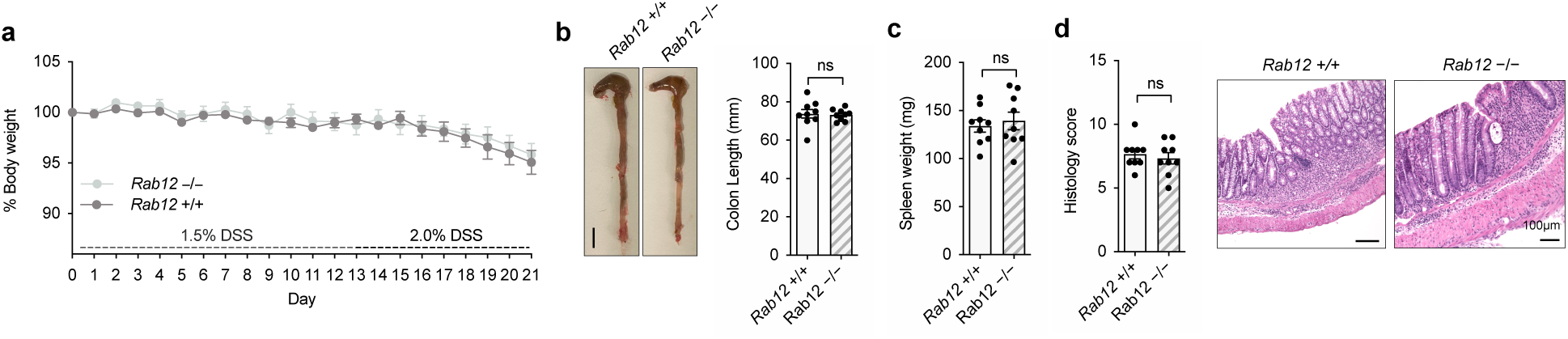
Assessment of WT and RAB12 KO Animals’ Response to DSS-Induced Colitis. **(a)** Normalized body weight of WT and RAB12 KO animals during progressively increasing DSS treatment over 21 days. **(b)** Representative colon photos and colon length measurements at sacrifice. **(c)** Spleen weight of WT and RAB12 KO animals at sacrifice. **(d)** Representative histology images and scoring (*n* = 18 animals).

## Methods and Materials

### Association analysis of LRRK2 variants with Crohn’s disease and Parkinson’s disease in three large biobanks

BioMe Biobank genotyping was performed using the Infinium Global Screening array (GSA, Illumina Inc.) and Infinium Global Diversity (GDA, Illumina Inc.) arrays. The GSA and GDA data were processed, QCed and imputed together using TOPMeD-r2 reference panel^28^. To determine genetic ancestries of Bio*Me* participants, the genotypes were combined with 1000 Genomes Project phase 3^29^ (1KGP) reference panel (https://www.cog-genomics.org/plink/2.0/resources, primary release, build 38, 3,202 samples). Cross-validation errors with K 4-12 were calculated using ADMIXTURE^30^, and GrafPOP^31^ was used to assign samples to a superpopulation. The Frobenius distance was used to compare each ADMIXTURE matrix K with the GrafPOP matrix, determining K=10 as the best fit for aligning self-identified and genetically determined matrices. The genetically determined classes have been unified to ancestry groups by identifying 1KGP reference samples with the highest ancestral proportions in each class.

Principal component analysis (PCA) was performed using linkage disequilibrium (LD)-pruned variants (*r*^2^ = 0.2) with a MAF greater than 5% and not exceeding HWE with a P < 1 × 10^−6^ using Plink v2^32^. CD and PD patients were identified based on electronic health records, containing ICD-10 and ICD-9 diagnoses of Bio*Me* participants matched with genotype data. CD patients were identified using K50 (ICD-10) and 555 (ICD-9) codes while PD patients were identified using G20 (ICD-10) and 332 (PD) codes.

Genebass (https://app.genebass.org/) provides exome-based association statistics of 394,841 UK BioBank participants^12^. We downloaded single-variant association statistics from gs://ukbb-exome-public/500k/results/variant_results.mt. Associations of the G2019, N2081D and N551K variants with CD (K50) and PD (G20) were extracted using Hail (https://github.com/hail-is/hail).

Million Veteran Program (MVP): Associations of the G2019, N2081D, and N551K variants with CD and PD were extracted from ancestry-specific and meta-analysis results available through the MVP PheWeb browser (https://phenomics.va.ornl.gov/web/cipher/pheweb)^33^.

### Association testing and meta-analysis of LRRK2 variants

Population-specific association testing was conducted in the Bio*Me* BioBank African American (AFR), Admixed American (AMR) and European American (EUR) cohorts using the SAIGE^34^. Analyses were adjusted for age, biological sex, array type and the first 10 principal components (PC 1-10) as covariates. For meta-analysis, we used METAL software^35^ to perform both European-specific and multi-ancestry meta-analyses using the inverse-variance weighted method.

### Plasmids

Dr. Christian Johannes Gloeckner (University of Tübingen, Germany) kindly provided the human LRRK2 pDEST-NSF-tandem affinity plasmid. We generated additional LRRK2 variants using the QuikChange Lightning Site-Directed Mutagenesis Kit as per the manufacturer’s instructions (Agilent, #210518).

### HEK Cell Culture, Transfection, and Lysis

HEK293 cells were cultured in DMEM (Dulbecco’s Modified Eagle Medium, Gibco™) containing 10% fetal calf serum, 100 U/ml penicillin, and 100 μg/ml streptomycin at 37°C in a humidified incubator with 5% CO2. Cells were seeded in six-well plates and transiently transfected at ∼75% confluency with 2 µg of DNA using Lipofectamine 2000, as per the manufacturer’s instructions (ThermoFisher Scientific). After 24 hours, cells were lysed in ice-cold 1x Cell Lysis Buffer (Cell Signaling #8903) supplemented with 1x protease and phosphatase inhibitors (ThermoFisher #78440) and left on ice for 30 minutes. Cell lysates were clarified by centrifugation at 15,000 g for 10 minutes at 4°C, and protein concentration was determined using the Pierce BCA Protein Assay Kit (ThermoFisher #23228). Fifteen micrograms of protein were loaded for quantitative immunoblot analysis.

### Structural analysis and graphic display

Structural model of LRRK2 (PDB ID: 7LI4) was manually examined using Coot^36^ and PyMol^37^ to glean the potential interactions occurring at N551 and N2081. The potential dynamics of these residues were analyzed using NAMD^38^. CHARMM force field was used to parameterize the simulation. The system was solvated inside a box of water molecules with a 13 Å padding in each direction. The system was then neutralized with 0.15 M NaCl. Simulation was performed with a 1 fs time step. The minimized structure was heated from 0 to 300 K over 300 ps. The production run was performed in the NVE (microcanonical) ensemble at 300 K. The total simulation time was 2 ns, and coordinates were recorded every 1 ps. Structural graphic presentations were made with PyMol.

### Generation of LRRK2 N2081D knock-in mouse model

The N2081D missense mutation was incorporated into the murine LRRK2 gene using gene editing (CRISPR/Cas9) technology as described previously (Shola et al., 2021). In brief, one gRNA (IDT: Integrated DNA Technologies Inc.) binding to the genomic target (GCTAACTCATCAAACTCATTGG) was selected after the evaluation of its off-target potential using the web-based tool CRISPOR (http://crispor.tefor.net) and its on-target efficiency by electroporating gRNA/Cas9 reagents (2 µM) into 0.5-day fertilized zygotes followed by sequence analysis of target genomic region of 2-cell stage embryos. Synthetic single-stranded editing template (IDT) contains the N2081D missense mutation, and flanking homology arms is as follows:

5’CTTACTACTTCACGATATTTGGACAACTGGGAGTAGGATTATGGAGGGTTTGAGGTTCCCAGATGAATTCGAT GAGTTAGCCATACAAGGGAAGTTGCCAGGTAAGTTCTGGTTTTATCTACAAGAGTTCTTTTCTTAATGTCA GCTTGGTCATGTAGAG.

Two silent mutations were also incorporated into the editing template to create a diagnostic EcoRI restriction enzyme recognition site and to prevent the recut by the gRNA/Cas9 DNA nuclease. The single-stranded donor DNA (10 ng/μL) was co-microinjected with pre-assembled gRNA/Cas9 riboproteins (0.2 µM of gRNA, 0.2µM of Cas9 protein, IDT) into the pronuclei of mouse zygotes in C57Black/6J background.

### Animals

The G2019S knock-in mouse line was kindly provided by Dianna Benson. RAB12KO mice were generated by our group in a previous study by deleting exon 3 of the *Rab12* gene in the C57BL/6J background using CRISPR-Cas9 technology. To generate N2081D with RAB12 KO mice, N2081D homozygotes were crossed with RAB12 KOs. These mice were then bred to produce RAB12 KO and N2081D homozygotes. All animal studies followed the guidelines of the Icahn School of Medicine at Mount Sinai Animal Care and Use Committee. Mice were housed under a 12-hour light-dark cycle.

### Tissue lysis

Organ tissue samples were collected, immediately frozen on dry ice, and stored at -80°C until lysis. Frozen tissues were homogenized in ice-cold 1x homogenization buffer (20 mM Tris-HCl, 0.5 mM EDTA, 250 mM sucrose) supplemented with 1x protease and phosphatase inhibitors (ThermoFisher #78440). Tissues were placed in screw-cap RINO microcentrifuge tubes containing stainless steel beads (NextAdvance) and homogenized using the Bullet Blender Gold (NextAdvance). The homogenate was combined with 10x CLB to achieve a 1x final concentration and rotated at 4°C for 30 minutes. Tissue lysates were centrifuged at 15,000 g for 10 minutes at 4°C, and supernatants were quantified using the Pierce BCA Protein Assay Kit (ThermoFisher #23228). For immunoblot analysis, 50 µg of protein per sample was loaded.

### Dextran Sulfate Sodium (DSS) Administration to Induce Colitis

To induce colitis, 2% (w/v) DSS (molecular weight 36,000-50,000 Da, MP Biomedicals) was dissolved in drinking water and provided ad libitum. The DSS solution was freshly prepared and replaced every 2-3 days for consistent dosing. For comparing DSS effects on WT, G2019S, and N2081D mice, the mice were 16 weeks old. For comparing N2081D mice and RAB12 KO effects, the mice were 12 weeks old. Mice whose body weight loss exceeded 20% of their original weight were euthanized as a humane endpoint.

Mice were monitored daily for signs of colitis, such as weight loss, stool consistency, and rectal bleeding. At the end of the DSS treatment or upon reaching humane endpoints, mice were euthanized, and colonic tissues were collected for histological and molecular analyses to assess colitis severity and inflammatory responses. Histological scoring was performed using a modified scale (excluding regeneration assessment) as described by Lamas et al. Stool samples were analyzed for Lipocalin-2 levels using ELISA (R&D Systems, #DY1857).

### Immunoblot analysis

Samples were mixed with 4× SDS–PAGE loading buffer supplemented with 150 mM DTT and heated at 95°C for 5 minutes. They were loaded onto NuPAGE 4–12% Bis–Tris Midi Gels (Thermo Fisher Scientific, Cat no. WG1402BOX or WG1403BOX) and electrophoresed at 120 V in NuPAGE MOPS SDS running buffer (Thermo Fisher Scientific, Cat no. NP0001-02). After electrophoresis, proteins were transferred onto a nitrocellulose membrane (GE Healthcare, Amersham Protran Supported 0.45 mm NC) at 20 V for 60 minutes in transfer buffer (25 mM Tris base, 190 mM glycine, and 15% methanol). The membranes were blocked with Intercept (TBS) blocking buffer (LicorBio) at room temperature for 1 hour.

They were incubated overnight at 4°C with the primary antibody in TBS blocking buffer mixed with TBS-T (50 mM Tris base, 150 mM NaCl, 0.1% Tween 20). Membranes were washed in TBS-T before incubation with the secondary antibody (IRDye, LicorBio) according to the manufacturer’s instructions. Following secondary antibody incubation, membranes were washed in TBS-T, and protein bands were detected using the Odyssey Classic Imaging System and quantified using Image Studio Lite.

### Culturing of bone marrow-dendritic cells (BMDCs)

Mouse femurs and tibias were flushed with a 23G needle using ice-cold PBS. Cells were pelleted by gentle centrifugation and resuspended in Red Blood Cell Lysis Buffer (eBioscience™ RBC Lysis Buffer, #00-4300-44) for 3 minutes, then diluted with PBS and strained through a 40-μm filter (Greiner EASYstrainer™, #542040). After pelleting, cells were resuspended in RPMI 1640 medium containing 10% fetal calf serum, 100 U/ml penicillin, and 100 μg/ml streptomycin, supplemented with 20 ng/ml GM-CSF (PeproTech) and 20 ng/ml IL-4 (PeproTech). On day 3, half of the medium was replaced. On day 6, cells were replated in supplemented RPMI medium and used for experiments on day 7. Cells were stimulated with RPMI medium containing Zymosan (InvivoGen) at 0.1 mg/ml for the indicated times. For confocal imaging, cells were treated with Zymosan A BioParticles™, Alexa Fluor™ 594 conjugate (ThermoFisher, #723374) and processed as described below.

### Phosphoproteomics and global proteomics sample preparation

Proteomics samples were first lysed with 100 µL of lysis buffer (60 mM Tetraethylammonium bromide (TEAB) pH 8.5, 5 mM Tris(2-carboxyethyl)phosphin (TCEP) and 25 mM 2-Chloroacetamide (CAA) in 10% Acetonitril (AcN)) for 30 min at 76 °C with agitation (1200 rpm) on an Eppendorf Thermomixer C. Afterwards, proteins were digested with Trypsin and LysC (Sigma Aldrich) with a 1:100 protein:enzyme ratio overnight at 37 °C and with agitation (1200 rpm). Formic acid was added to a final concentration of 1% to stop the digestion. Samples were placed in a speedvac at 30 °C and spun under vacuum until full dryness. Samples were reconstituted in 0.1% Formic acid. For global proteomics, 200 ng of peptides were subjected to Evotips purification, see below. For phospho-peptide enrichment 200 µg of peptides in 100 µL were subjected to enrichment using the default phospho-peptide enrichment protocol from the AssayMAP Bravo Platform (Agilent). For the enrichment 50 ml of 0.1% formic acid (FA) in AcN was used as the priming buffer, 50 ml 0.1% FA in 80 % AcN as the equilibration buffer and 50 ml of 500 mM ammonium hydrogen phosphate as the elution buffer with Fe(III)-NTA cartridges. For further information we refer to: https://www.agilent.com/cs/library/flyers/public/5991-6101EN.pdf (Evosep). Evotips were first soaked in 1-propanol for 3 minutes, then washed two times with 0.1% formic acid in 100% acetonitrile (EvoB), soaked for 3 minutes in 1-propanol and then washed again two times with 0.1% formic acid in water (EvoA) all at 700 g for 1 min. 70 µL of EvoA were loaded on Evotips at 700 g for 15 seconds before sample were loaded at 700 g for 1 min. Evotips were washed with 50 µL EvoA. Additional 150 µL of EvoA was loaded on top at 700 g for 15 seconds. Samples were then subjected to LC-MS analysis.

### Data-independent acquisition LC-MS analysis

For the global proteomics experiment, samples were subjected to LC-MS/MS analysis on a Orbitrap Astral mass spectrometer (Thermo Fisher Scientific) couple online to an Evosep One liquid chromatography (LC) system (Evosep). Peptides were eluted from the Evotips with up to 35% ACN and separated using a 21 min gradient for a throughput of 60 samples per day (SPD) on an Aurora Rapid TS column of 8 cm, 150 μm i.d. with 1.7 μm C18-beads (IonOpticks). Column temperature was maintained at 50 °C using a column oven (IonOpticks). The Orbitrap Astral MS was equipped with a FAIMS Pro interface and runs were acquired using a FAIMS compensation voltage of -40 V and a total carrier gas flow of 3.5 L/min. Full MS scans was in the range of 380 - 980 m/z with an Orbitrap resolution of 120,000 with a normalized automated gain control (AGC) of 500% and a maximum injection time of 3 ms. For Astral MS/MS scans in data-independent mode, the isolation windows were set to 3 Th with a maximum injection time of 5 ms and an AGC target of 800%. Isolated ions were fragmented using HCD with a normalized collision energy of 25%.

For phospho-proteomics analysis, we used a timsTOF Pro 2 mass spectrometer (Bruker) coupled to the Evosep One system. Peptides were eluted from Evotips with up to 35% ACN and separated using a 44 min gradient for 30 SPD on a PepSep C18 column (15 cm x 150 µm, 1.5 µm, Bruker). The dia-PASEF method consisted of eight dia-PASEF scans, which incorporated two ion mobility (IM) windows per dia-PASEF scan (cycle time 2.7 seconds). The method covered a mass-to-charge (m/z) range of 400 to 1400 and an ion mobility range from 0.75 to 1.45 Vs cm−2. For all experiments, we used accumulation and ramp times of 100 ms. We implemented a constantly decreasing collision energy profile from 60 eV at 1.5 Vs cm−2 to 54 eV at 1.17 Vs cm−2 to 25 eV at 0.85 Vs cm−2 and end at 20 eV at 0.6 Vs cm−2.

### Bioinformatics analysis

For the global proteomics experiment DIA raw files were processed using the library free search in DIA-NN 1.8.1 (DIA-NN: Demichev et al, Nature Methods, 2020, https://www.nature.com/articles/s41592-019-0638-x) against a UniProt mouse reference proteome of canonical sequences with 17214 entries, where the enzyme specificity was set to trypsin, with a maximum number of one missed cleavage site, a maximum peptide length of 30 and minimum of 7, up to one variable modification, carbamidomethylation as a fixed modifcation, while oxidation of methionine was set as a variable modification. The deep learning based spectra mode, FASTA digest for library free search and RTs and IMs prediction with heuristic protein inference was enabled. FDR control for precursor was set to 1% while the remaining settings were set to default.

For the phosphopeptide experiment, DIA raw files were processed using directDIA in Spectronaut against the same Uniprot mouse reference proteome. Default settings were used with the following exceptions: variable Modification were set to Phospho (STY) with 3-25 best fragments per peptide, normalization filter type set to phospho (STY) and a probability cutoff at 0. Peptide table was retrieved in a suitable format for the peptide collaps plugin described in Perseus (https://www.nature.com/articles/nmeth.3901). Peptide collapse was done with default setting except for the aggregation type, which was set to sum.

For the bioinformatics analysis Python version 3.5.5 with pandas 1.4.2, numpy 1.21.5, matplotlib 3.5.13, seaborn 0.11.2, scipy 1.7.3 and statsmodels 0.13.2 were used. Protein and peptide intensities were first normalised using directLFQ (citations) and then log_2_-transformed. Data was filtered for valid values in at least one experimental group and imputed using a sampleing method with a shifted Gaussian normal distribution (width = 0.3 and downshift = 1.8). Statistical significance was determined using an unpaired two-sided student’s t-test. P-value correction was done using the Benjamini-Hochberg method.

### Antibodies

The antibodies used for immunoblot analysis were: Mouse monoclonal antibody to FLAG peptide (Sigma #F1804, 1:5000), Mouse monoclonal antibody to β-actin (Cell Signaling #4967, 1:5000), Rabbit monoclonal antibody to LRRK2 (Abcam #MJFF2, ab133474, 1:2000), Rabbit monoclonal antibody to LRRK2 phospho-S1292 (Abcam #ab203181, 1:1000), Rabbit monoclonal antibody to total-RAB10 (Abcam #ab237703, 1:2000), and Rabbit monoclonal antibody to RAB10 phospho-T73 (Abcam #ab230261, 1:1000). For immunocytochemistry, the antibodies used were: Rabbit monoclonal antibody to RAB10 phospho-T73 (Abcam #ab230261, 1:100), and Rabbit monoclonal antibody to LRRK2 (Abcam #MJFF2, ab133474, 1:100).

### Confocal microscopy

Cells were seeded on coverslips and washed with PBS-T. They were fixed in 4% paraformaldehyde in PBS for 15 minutes at room temperature. The cells were blocked with 1% BSA (Sigma-Aldrich) in PBS plus 0.1% Triton X-100 for 30 minutes. Primary antibodies were diluted in 1% BSA in PBS plus 0.1% Triton X-100 and incubated at 4°C overnight. After washing three times with PBS-T, the cells were incubated with secondary antibodies for 60 minutes at room temperature in the dark. The coverslips were mounted on glass slides with ProLong Gold Antifade Mountant with DAPI (Life Technologies). Slides were imaged using an LSM 900 microscope.

### Statistical analysis

Statistical differences in HEK293 cell experiments were calculated using one-way ANOVA with Dunnett’s multiple comparison test, comparing each group to WT. Comparisons between WT, G2019S, and N2081D samples were conducted using one-way ANOVA with Tukey’s post hoc test for multiple comparisons. For comparisons between two experimental groups, an unpaired two tailed t-test was used.

## Acknowledgments

This work was supported in part through the computational resources and staff expertise provided by Scientific Computing and Data at the Icahn School of Medicine at Mount Sinai and supported by the Clinical and Translational Science Awards (CTSA) grant UL1TR004419 from the National Center for Advancing Translational Sciences. We would also like to thank Mabel Ko for her training and consultation on histological scoring. Our gratitude extends to Ji Sun for providing the structural models of LRRK2. Additionally, we thank Meriem Belabed for her assistance in optimizing our BMDC culture conditions.

## References

1 Healy, D. G. et al. Phenotype, genotype, and worldwide genetic penetrance of LRRK2-associated Parkinson’s disease: a case-control study. Lancet Neurol 7, 583–590 (2008). 10.1016/s1474-4422(08)70117-0

2 Bloem, B. R., Okun, M. S. & Klein, C. Parkinson’s disease. Lancet 397, 2284–2303 (2021). 10.1016/s0140-6736(21)00218-x

3 Hui, K. Y. et al. Functional variants in the LRRK2 gene confer shared effects on risk for Crohn’s disease and Parkinson’s disease. Sci Transl Med 10 (2018). 10.1126/scitranslmed.aai7795

4 Alessi, D. R. & Sammler, E. LRRK2 kinase in Parkinson’s disease. Science 360, 36–37 (2018). doi:10.1126/science.aar5683

5 Steger, M. et al. Phosphoproteomics reveals that Parkinson’s disease kinase LRRK2 regulates a subset of Rab GTPases. eLife 5, e12813 (2016). 10.7554/eLife.12813

6 Berwick, D. C., Heaton, G. R., Azeggagh, S. & Harvey, K. LRRK2 Biology from structure to dysfunction: research progresses, but the themes remain the same. Mol Neurodegener 14, 49 (2019). 10.1186/s13024-019-0344-2

7 Liu, Z., Xu, E., Zhao, H. T., Cole, T. & West, A. B. LRRK2 and Rab10 coordinate macropinocytosis to mediate immunological responses in phagocytes. Embo j 39, e104862 (2020). 10.15252/embj.2020104862

8 Herbst, S. et al. LRRK2 activation controls the repair of damaged endomembranes in macrophages. Embo j 39, e104494 (2020). 10.15252/embj.2020104494

9 Bonet-Ponce, L. et al. LRRK2 mediates tubulation and vesicle sorting from lysosomes. Sci Adv 6 (2020). 10.1126/sciadv.abb2454

10 Kluss, J. H., Bonet-Ponce, L., Lewis, P. A. & Cookson, M. R. Directing LRRK2 to membranes of the endolysosomal pathway triggers RAB phosphorylation and JIP4 recruitment. Neurobiol Dis 170, 105769 (2022). 10.1016/j.nbd.2022.105769

11 Dolinger, M., Torres, J. & Vermeire, S. Crohn’s disease. Lancet 403, 1177–1191 (2024). 10.1016/s0140-6736(23)02586-2

12 Karczewski, K. J. et al. Systematic single-variant and gene-based association testing of thousands of phenotypes in 394,841 UK Biobank exomes. Cell Genom 2, 100168 (2022). 10.1016/j.xgen.2022.100168

13 Verma, A. et al. Diversity and scale: Genetic architecture of 2068 traits in the VA Million Veteran Program. Science 385, eadj1182 (2024). doi:10.1126/science.adj1182

14 Kars, M. E. et al. The landscape of rare genetic variation associated with inflammatory bowel disease and Parkinson’s disease comorbidity. Genome Medicine 16, 66 (2024). 10.1186/s13073-024-01335-2

15 Matikainen-Ankney, B. A. et al. Altered Development of Synapse Structure and Function in Striatum Caused by Parkinson’s Disease-Linked LRRK2-G2019S Mutation. J Neurosci 36, 7128–7141 (2016). 10.1523/jneurosci.3314-15.2016

16 Wirtz, S. et al. Chemically induced mouse models of acute and chronic intestinal inflammation. Nat Protoc 12, 1295–1309 (2017). 10.1038/nprot.2017.044

17 Rutella, S. & Locatelli, F. Intestinal dendritic cells in the pathogenesis of inflammatory bowel disease. World J Gastroenterol 17, 3761–3775 (2011). 10.3748/wjg.v17.i33.3761

18 Takagawa, T. et al. An increase in LRRK2 suppresses autophagy and enhances Dectin-1-induced immunity in a mouse model of colitis. Sci Transl Med 10 (2018). 10.1126/scitranslmed.aan8162

19 Berndt, B. E., Zhang, M., Chen, G. H., Huffnagle, G. B. & Kao, J. Y. The role of dendritic cells in the development of acute dextran sulfate sodium colitis. J Immunol 179, 6255–6262 (2007). 10.4049/jimmunol.179.9.6255

20 Kubo, M. et al. Leucine-Rich Repeat Kinase 2 Controls Inflammatory Cytokines Production through NF-κB Phosphorylation and Antigen Presentation in Bone Marrow-Derived Dendritic Cells. Int J Mol Sci 21 (2020). 10.3390/ijms21051890

21 Liu, Z. et al. The kinase LRRK2 is a regulator of the transcription factor NFAT that modulates the severity of inflammatory bowel disease. Nature Immunology 12, 1063–1070 (2011). 10.1038/ni.2113

22 Kluss, J. H. et al. Preclinical modeling of chronic inhibition of the Parkinson’s disease associated kinase LRRK2 reveals altered function of the endolysosomal system in vivo. Mol Neurodegener 16, 17 (2021). 10.1186/s13024-021-00441-8

23 Fell, M. J. et al. MLi-2, a Potent, Selective, and Centrally Active Compound for Exploring the Therapeutic Potential and Safety of LRRK2 Kinase Inhibition. Journal of Pharmacology and Experimental Therapeutics 355, 397–409 (2015). 10.1124/jpet.115.227587

24 Dhekne, H. S. et al. Genome-wide screen reveals Rab12 GTPase as a critical activator of Parkinson’s disease-linked LRRK2 kinase. Elife 12 (2023). 10.7554/eLife.87098

25 Wang, X. et al. Rab12 is a regulator of LRRK2 and its activation by damaged lysosomes. eLife 12, e87255 (2023). 10.7554/eLife.87255

26 Li, X. et al. RAB12-LRRK2 Complex Suppresses Primary Ciliogenesis and Regulates Centrosome Homeostasis in Astrocytes. bioRxiv, 2024.2007.2017.603999 (2024). 10.1101/2024.07.17.603999

27 Cabezudo, D., Tsafaras, G., Van Acker, E., Van den Haute, C. & Baekelandt, V. Mutant LRRK2 exacerbates immune response and neurodegeneration in a chronic model of experimental colitis. Acta Neuropathologica 146, 245–261 (2023). 10.1007/s00401-023-02595-9

28 Taliun, D. et al. Sequencing of 53,831 diverse genomes from the NHLBI TOPMed Program. Nature 590, 290–299 (2021). 10.1038/s41586-021-03205-y

29 Auton, A. et al. A global reference for human genetic variation. Nature 526, 68–74 (2015). 10.1038/nature15393

30 Alexander, D. H., Novembre, J. & Lange, K. Fast model-based estimation of ancestry in unrelated individuals. Genome Res 19, 1655–1664 (2009). 10.1101/gr.094052.109

31 Jin, Y., Schaffer, A. A., Feolo, M., Holmes, J. B. & Kattman, B. L. GRAF-pop: A Fast Distance-Based Method To Infer Subject Ancestry from Multiple Genotype Datasets Without Principal Components Analysis. G3 (Bethesda) 9, 2447–2461 (2019). 10.1534/g3.118.200925

32 Purcell, S. et al. PLINK: a tool set for whole-genome association and population-based linkage analyses. Am J Hum Genet 81, 559–575 (2007). 10.1086/519795

33 Verma, A. et al. Diversity and scale: Genetic architecture of 2068 traits in the VA Million Veteran Program. Science 385, eadj1182 (2024). 10.1126/science.adj1182

34 Zhou, W. et al. Efficiently controlling for case-control imbalance and sample relatedness in large-scale genetic association studies. Nat Genet 50, 1335–1341 (2018). 10.1038/s41588-018-0184-y

35 Willer, C. J., Li, Y. & Abecasis, G. R. METAL: fast and efficient meta-analysis of genomewide association scans. Bioinformatics 26, 2190–2191 (2010). 10.1093/bioinformatics/btq340

36 Emsley, P. & Cowtan, K. Coot: model-building tools for molecular graphics. Acta Crystallogr D Biol Crystallogr 60, 2126–2132 (2004). https://doi.org/S0907444904019158 10.1107/S0907444904019158

37 The PyMOL Molecular Graphics System, V. S., LLC.

38 Phillips, J. C. et al. Scalable molecular dynamics on CPU and GPU architectures with NAMD. J Chem Phys 153, 044130 (2020). 10.1063/5.0014475

